# Roles of the EnvZ/OmpR Two-Component System and Porins in Iron Acquisition in *Escherichia coli*

**DOI:** 10.1101/2020.05.11.089920

**Authors:** Henri Gerken, Phu Vuong, Ketaki Soparkar, Rajeev Misra

## Abstract

*Escherichia coli* secretes high-affinity Fe^3+^ chelators to solubilize and transport chelated Fe^3+^ via specific outer membrane receptors. In microaerobic and anaerobic growth environments, where the reduced Fe^2+^ form is predominant, ferrous transport systems fulfill the bacterial need for iron. Expression of genes coding for iron metabolism is controlled by Fur, which when bound to Fe^2+^ acts as a repressor. Work carried out in this paper shows that the constitutively activated EnvZ/OmpR two-component system, which normally controls expression of the *ompC* and *ompF* porin genes, dramatically increases the intracellular pool of accessible iron, as determined by whole-cell electron paramagnetic resonance (EPR) spectroscopy, by inducing the OmpC/FeoB-mediated ferrous transport pathway. Elevated levels of intracellular iron in turn activated Fur, which inhibited the ferric but not the ferrous transport pathway. The data show that the positive effect of constitutively activated EnvZ/OmpR on *feoB* expression is sufficient to overcome the negative effect of activated Fur on *feoB*. In a *tonB* mutant, which lacks functional ferric transport systems, deletion of *ompR* severely impairs growth on rich medium not supplemented with iron, while the simultaneous deletion of *ompC* and *ompF* is not viable. These data, together with the observation of de-repression of the Fur regulon in an OmpC mutant, show that the porins play an important role in iron homeostasis. The work presented here also resolves a long-standing paradoxical observation of the effect of certain mutant *envZ* alleles on iron regulon.

**IMPORTANCE:** The work presented here solved a long-standing paradox of the negative effects of certain missense alleles of *envZ*, which codes for kinase of the EnvZ/OmpR two component system, on the expression of ferric uptake genes. The data revealed that the constitutive *envZ* alleles activate the Feo- and OmpC-mediated ferrous uptake pathway to flood the cytoplasm with accessible ferrous iron. This activates the ferric uptake regulator, Fur, which inhibits ferric uptake system but cannot inhibit the *feo* operon due to the positive effect of activated EnvZ/OmpR. The data also revealed importance of porins in iron homeostasis.

## INTRODUCTION

Iron, used as a redox center by many enzymes, is an essential trace metal required by almost all living organisms. The intracellular level of free catalytically active iron is typically kept low due to its toxic effects. Free ferrous iron reacts with hydrogen peroxide, a natural byproduct of aerobic respiration, to generate highly toxic hydroxyl radicals (OH^•^) via the Fenton reaction (1). Due to this potentially damaging property of iron, there exists an intricate balance between iron transport, utilization and storage. Most bacteria possess mechanisms to import iron in its oxidized ferric state (Fe^3+^), reduced ferrous state (Fe^2+^) or both (for reviews, see 2, 3). The solubility of these two iron forms differs drastically at neutral pH: ferric iron has extremely low solubility at 10^−18^ M, whereas ferrous iron is readily soluble at 10^−1^ M. To take up ferric iron, bacteria have developed high-affinity ferric iron chelators called siderophores to capture, solubilize, and deliver insoluble iron into the cell (4). Unlike the Fe^3+^ transport system, which requires a number of proteins involved in siderophore synthesis and Fe^3+^-siderophore acquisition, the Fe^2+^ transport system appears to consist of mainly one protein, FeoB (5). The FeoB protein is synthesized from the *feoABC* operon, whose expression is activated by Fnr, an anaerobic transcriptional regulator (5). FeoB is a highly conserved, 773-residue inner membrane protein that contains several GTP-binding motifs (6, 7, 8). In the absence of FeoB or FeoA, Fe^2+^ uptake is either virtually abolished (Δ*feoB*) or mildly reduced (Δ*feoA*) (5). The function of FeoC, which is present only in members of the *Enterobacteriaceae* family, is unknown (7, 8). FeoB and its homologs are required for full virulence in many bacteria, including *E. coli* (9), *Salmonella* Typhimurium (10, 11), and *Helicobacter pylori* (12).

Fur (<u>f</u>erric <u>u</u>ptake <u>r</u>egulator) in *E. coli* and its orthologs in many Gram-negative and Gram-positive bacteria are the master regulator of genes encoding both ferric and ferrous iron acquisition functions, as well as siderophore synthesis and uptake (13, 14). Cells lacking Fur experience iron overload that causes oxidative damage and mutagenesis (15). Fur-regulated genes contain one or more Fur-binding sites around the -35 and -10 regions of the promoter, often referred to as the Fur boxes (16, 17). Fur uses Fe^2+^ as a co-factor: when the level of available Fe^2+^ increases in the cell, it binds to Fur and enhances its affinity for DNA by almost 1,000-fold (2). The active Fur-Fe^2+^ complex then binds to a Fur box and represses transcription of the iron acquisition gene. RyhB is a small regulatory RNA whose transcription is also repressed by Fur-Fe^2+^ (18). Consequently, when Fur is active, the levels of RyhB are low, resulting in stabilization and translation of over a dozen mRNAs encoding non-essential iron-utilization proteins, including those that store iron (Bfr), detoxify superoxide (SodB), and catalyze steps of the TCA cycle (AcnA and SdhCDAB) (19). Thus, excess Fe^2+^ activates Fur to halt further iron uptake and at the same time, promotes the utilization of Fe^2+^, and inversely, low intracellular iron level induces iron uptake and utilization (20). Recent genome-wide analyses revealed a more comprehensive profile of Fur and RyhB regulons (21, 22).

Whereas Fur and RyhB are the principal determinants of iron homeostasis in *E. coli*, evidence exists supporting the involvement of some two-component signal transduction systems (TCSs) in iron homeostasis. EnvZ and OmpR are the archetypal TCS in *E. coli*, where EnvZ serves as a sensor kinase and OmpR as a response regulator (23). They respond to medium osmolarity and influence the expression of OmpC and OmpF, the two major porins that facilitate the diffusion of small hydrophilic solutes (∼600 Da) across the outer membrane (24). OmpC is preferentially expressed in high osmolarity, whereas OmpF expression is favored in low osmolarity (25). Microarray data from an Δ*ompR* Δ*envZ* background showed a significant increase in the expression of a number of Fur-regulated genes, particularly those involved in enterobactin siderophore synthesis and transport (26). Over three decades ago, Lundrigan and Earhart (27) reported that in a *perA* (*envZ*) mutant background, the levels of three iron-regulated proteins were significantly reduced. The authors suggested that this could be due to a posttranscriptional defect. Later, it was speculated that this inhibition could be due to the indirect effects of *envZ*/*ompR*, leading to alterations in the structure and diffusion properties of the outer membrane (28). While characterizing revertants of an *E. coli* mutant defective in outer membrane biogenesis, we discovered several pleiotropic *envZ* alleles conferring an OmpC^+^ OmpF^−^ LamB^−^ phenotype (29). These alleles were hypothesized to biochemically lock EnvZ into a conformation that causes increased OmpR phosphorylation. This activated EnvZ/OmpR state is thought to enable OmpR to bind to promoters with weak OmpR-binding affinities. One such pleiotropic *envZ* allele, *envZ*_*R397L*_, was characterized in detail (29). The preliminary whole genomic microarray analysis of the *envZ*_*R397L*_ mutant carried out in our laboratory found that the largest group of genes (>50) affected by the activated EnvZ_R397L_/OmpR^+^ background belonged to the Fur regulon (30; Table S1).

In this study, we show that EnvZ_R397L_ exerts its effect on the Fur regulon in part by increasing the accessible intracellular pool of iron via the OmpC-FeoB-mediated Fe^2+^ transport pathway. This, in turn, activates Fur and downregulates the Fe^3+^ transport pathway. Our analyses also revealed the critical roles of EnvZ/OmpR and porins in iron homeostasis in the Δ*tonB* background where high-affinity iron transport systems are non-functional.

## RESULTS

### Effects of *envZ*_*R397L*_ on the ferric transport system

We first set out to investigate the effects of *envZ*_*R397L*_ on the Fur regulon. RNA isolated from mid-log phase grown cells was converted to cDNA and levels of various transcripts were analyzed by RT-qPCR. The data in Fig. 1 show relative transcript levels of four Fur-regulated genes: *fecA, fepA, fhuA* and *fhuF*. In the *envZ*_*R397L*_ background, their transcript levels went down 10 (*fecA*), 3 (*fhuA*), and 2.5 (*fepA* and *fhuF*) folds relative to the wild type (EnvZ^+^) strain. As expected, in a Δ*fur* background their expression was de-repressed, resulting in a dramatic increase in their transcripts (Fig. 1). In that background, the presence of *envZ*_*R397L*_ was still able to reduce *fecA* and *fepA* transcript levels 3.6 and 9.5 folds, respectively, but not that of *fhuA* and *fhuF*, which experienced less than 20% reduction (Fig. 1). Using the chromosomally integrated *fepA*::*lacZ* and *fhuA*::*lacZ* gene fusion constructs, we were able to recapitulate the key RNA data shown in Fig. 1 (Fig. S1). This indicated that EnvZ_R397L_/OmpR or factors under the activated TCS’s control could also down-regulate *fecA* and *fepA* transcription in the absence of Fur. In contrast, the negative effect of *envZ*_*R397L*_ on *fhuA* and *fhuF* expression requires Fur. Moreover, the repressive effect of *envZ*_*R397L*_ on *fecA* and *fepA* in the *fur*^+^ background was found to be independent of OmrA and OmrB (Fig. S2), the two EnvZ/OmpR-dependent small regulatory RNAs whose overexpression from plasmids was previously shown to down-regulate *fecA, fepA* and other Fur-regulated genes (31). It is worth mentioning that the *envZ*_*R397L*_ allele has been previously shown to increase *omr*::*lacZ* expression almost tenfold (29).

**Fig. 1.**
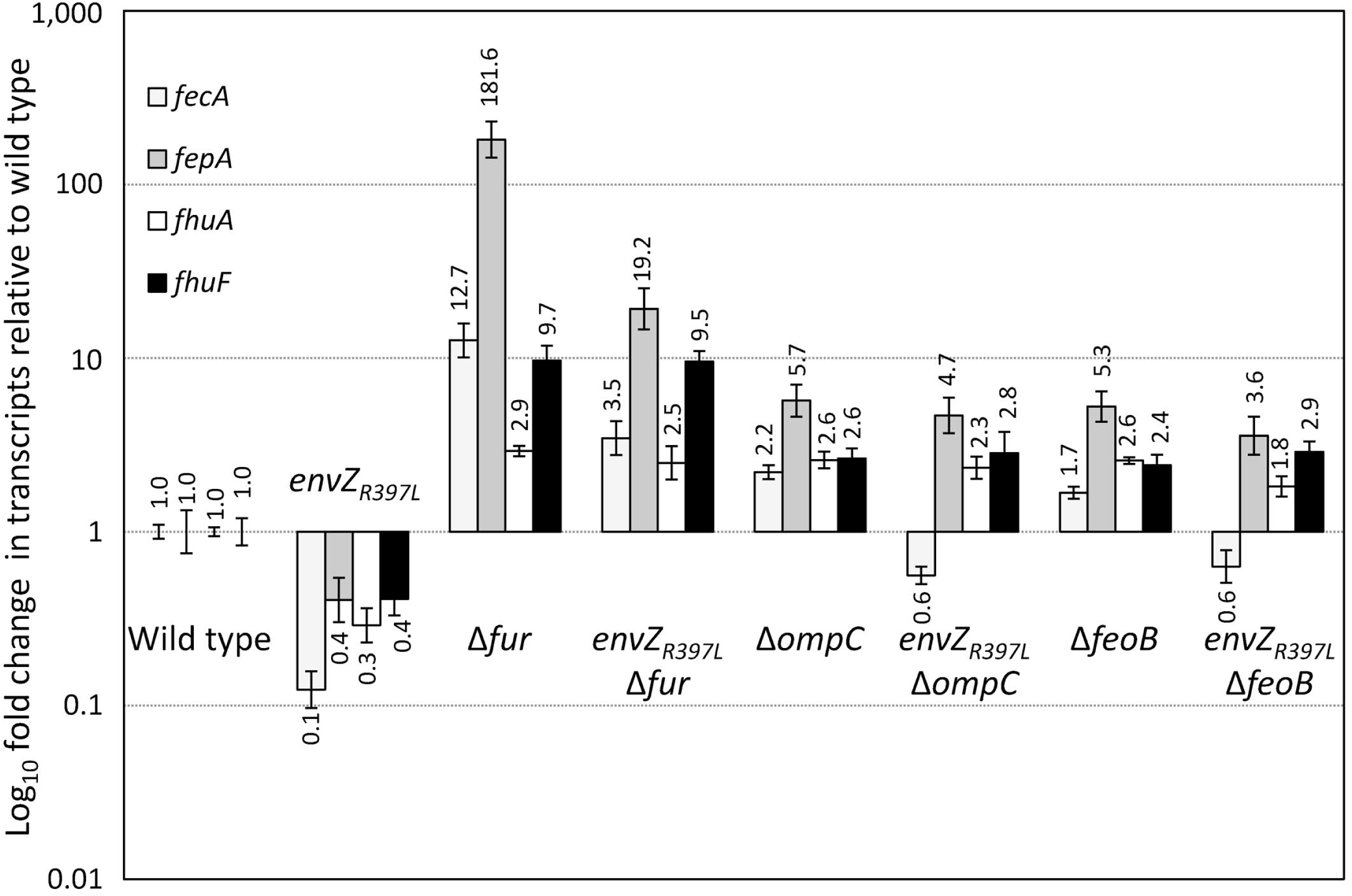
Determination of *fecA, fepA fhuA* and *fhuF* expression in different genetic backgrounds by real-time quantitative PCR (RT-qPCR). RNA was isolated from bacterial cultures grown to mid-log. Relative quantification of transcripts was performed using the 2^-ΔΔCT^ method, with *ftsL* and *purC* serving as the reference genes. Relative fold-changes in gene expression and error bars representing standard deviation are shown. Bacterial strains used are: RAM1292 (wild type), RAM1541 (*envZ*_*R397L*_), RAM2697 (Δ*fur*), RAM2698 (*envZ*_*R397L*_ Δ*fur*), RAM2699 (Δ*ompC*), RAM2700 (*envZ*_*R397L*_ Δ*ompC*), RAM2701 (Δ*feoB*), and RAM2702 (*envZ*_*R397L*_ Δ*feoB*).

Expression of OmpC is activated constitutively in the *envZ*_*R397L*_ background, while that of OmpF and LamB is severely inhibited (29). To ask whether OmpC is somehow involved in the *envZ*_*R397L*_-mediated down-regulation of *fecA, fepA, fhuA* and *fhuF*, we examined their transcript levels in the Δ*ompC* and Δ*ompC envZ*_*R397L*_ backgrounds. Remarkably, without OmpC, *envZ*_*R397L*_ was unable to exert any significantly negative effect on *fepA, fhuA* and *fhuF* expression, while the effect on *fecA* diminished from tenfold in the presence of OmpC to less than twofold without OmpC (Fig. 1). Interestingly, the levels of all four transcripts went up in Δ*ompC* cells (Fig. 1). We theorize that without OmpC, diffusion of Fe^2+^ into the cell is decreased and the less active Fur fails to fully repress *fecA, fepA, fhuA* and *fhuF* expression.

If the intake of Fe^2+^ by OmpC porin increases active Fur-Fe^2+^ levels, then the absence of FeoB, the Fe^2+^-specific iron transporter, should also interfere with this activation and abrogate the Fur-mediated effects of *envZ*_*R397L*_ on *fecA, fepA, fhuA* and *fhuF*. Indeed, just like in the Δ*ompC* background, *fecA, fepA, fhuA* and *fhuF* transcript levels went up in the Δ*feoB* background, and *envZ*_*R397L*_ could either no longer impose a significant negative effect (*fepA, fhuA* and *fhuF*) or the effect was significantly reduced (*fecA*).

### Effects of *envZ*_*R397L*_ on the ferrous transport system

The data presented in Fig. 1 showed the involvement of the FeoB ferrous iron transporter and OmpC porin in *envZ*_*R397L*_-mediated down-regulation of the ferric iron transport system. While *ompC* expression increases in the *envZ*_*R397L*_ background (29), the status of the *feo* operon in this background is unknown. The *feo* operon is under the negative control of Fur (5). Consequently, if higher Fur-Fe^2+^ activity is present in the *envZ*_*R397L*_ background, as we have suggested above, then the expression of the *feo* operon, like that of *fecA, fepA, fhuA* and *fhuF*, should also be inhibited. This, however, will be incongruent with our data showing *envZ*_*R397L*_’s dependence on *feoB* for its effects. We therefore hypothesized that *feo* expression, like that of *ompC*, is activated by *envZ*_*R397L*_ to such a degree that it more than compensated for the *feo* down-regulation by increased Fur-Fe^2+^ activity.

To test these possibilities, we analyzed *feoA* and *feoB* transcript levels in different genetic backgrounds by RT-qPCR (Fig. 2). Note that *feoABC* are part of a contiguous operon and therefore, likely expressed from a polycistronic message. Consequently, *feoA* and *feoB* transcript analysis probes their respective coding regions in a polycistronic message. In the EnvZ_R397L_ background, *feoA* and *feoB* transcript levels went up dramatically over those in the *envZ*^+^ control strain. As expected, their levels also went up in the Δ*fur* background. Interestingly, in the *envZ*_*R397L*_ Δ*fur* background *feoA* and *feoB* transcript levels increased well above those in the individual mutation backgrounds, indicating that *envZ*_*R397L*_ and Δ*fur* act independently and synergistically to enhance *feo* expression. Again these observations were recapitulated using the chromosomally integrated *feo*::*lacZ* fusion (Fig. S1). These data support our hypothesis that *envZ*_*R397L*_ activates *feo* expression in a fashion that counteracts repression by higher levels of Fur-Fe^2+^.

**Fig. 2.**
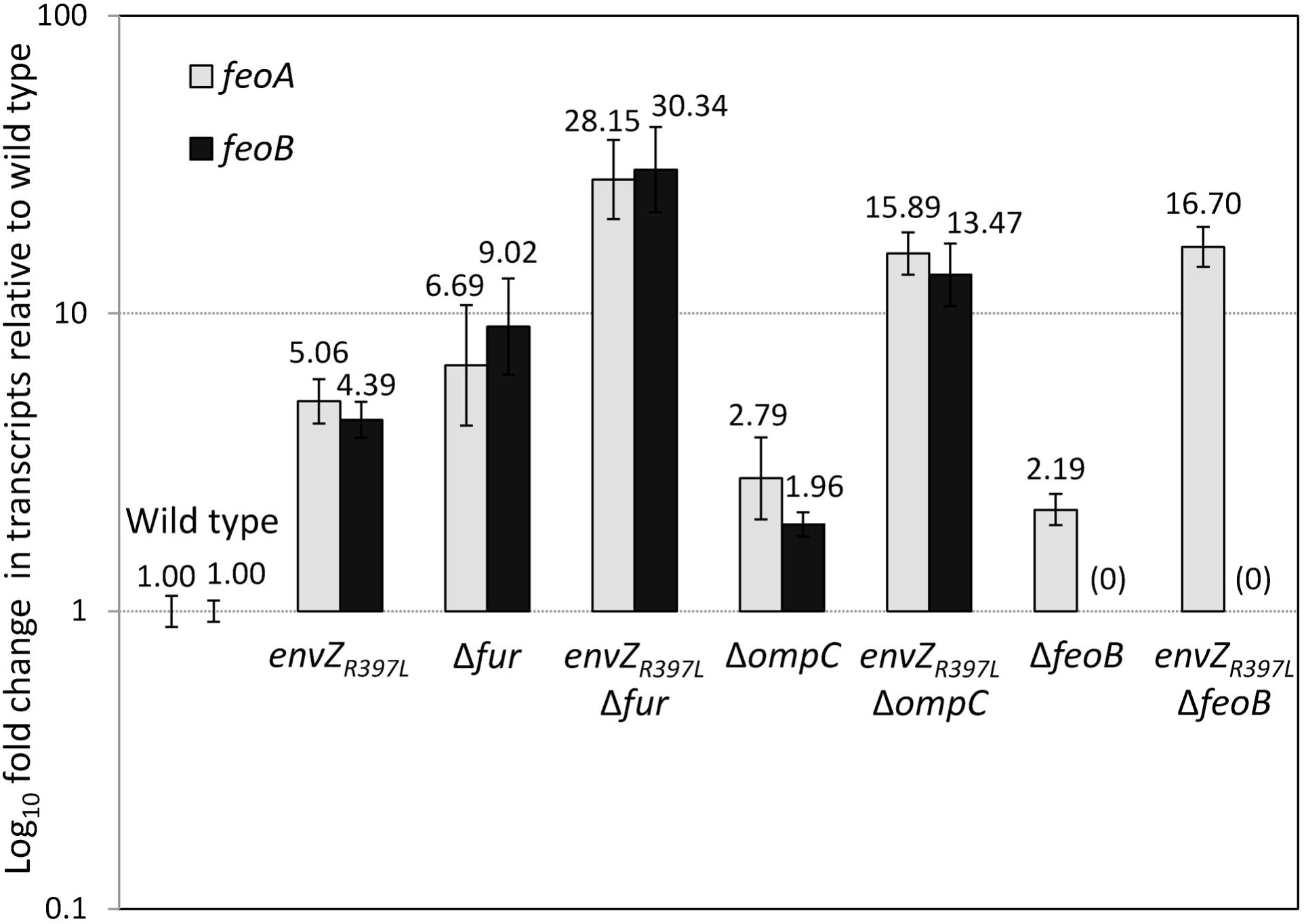
Determination of the relative gene expression of *feoA* and *feoB* by RT-qPCR. RNA was isolated from bacterial cultures grown to mid-log. Relative quantification of transcripts in various genetic backgrounds was performed using the 2^-ΔΔCT^ method, with *ftsL* and *purC* serving as reference genes. Relative fold-changes in gene expression and error bars representing standard deviation are shown. Bacterial strains used are: RAM1292 (wild type), RAM1541 (*envZ*_*R397L*_), RAM2697 (Δ*fur*), RAM2698 (*envZ*_*R397L*_ Δ*fur*), RAM2699 (Δ*ompC*), RAM2700 (*envZ*_*R397L*_ Δ*ompC*), RAM2701 (Δ*feoB*), and RAM2702 (*envZ*_*R397L*_ Δ*feoB*).

We then examined the effects of *envZ*_*R397L*_ on *feoA* and *feoB* transcript levels in the absence of OmpC or FeoB. Without OmpC or FeoB, a modest twofold increase in *feoA* and *feoB* (Δ*ompC*) or *feoA* (Δ*feoB*) transcripts was observed (Fig. 2). We interpret this to reflect a modest relief in the Fur-mediated repression of the *feo* operon, since we have already implicated OmpC and FeoB in the ferrous iron transport and increase in Fur-Fe^2+^ levels (Fig. 1). The presence of *envZ*_*R397L*_ in the Δ*ompC* or Δ*feoB* background led to an increase in *feoA* and *feoB*, or *feoA* transcripts, respectively, in a synergistic fashion, which is likely due to the simultaneous activation of *feo* expression by *envZ*_*R397L*_ and a modest decrease in the Fur-mediated repression of *feo* from the absence of OmpC and FeoB. These data showed that *envZ*_*R397L*_ inhibits ferric transport pathway but activates ferrous transport pathway.

### Intracellular iron levels in the *envZ*_*R497L*_ mutant

The OmpC/Feo-mediated increase in Fur-Fe^2+^ activity in the *envZ*_*R397L*_ background implies that the cytoplasm of the *envZ*_*R397L*_ mutant contains higher levels of accessible iron than that in the cytoplasm of the EnvZ^+^ cell. To test this directly, we measured the intracellular pool of accessible iron by whole-cell electron paramagnetic resonance (EPR) spectroscopy, a method established in the Imlay laboratory (32). The data presented in Fig. 3 show that the wild type (EnvZ^+^) strain had 32 μM of accessible intracellular iron. Expectedly, this level rose fourfold to 120 μM in the Δ*fur* mutant. Remarkably, the level of accessible iron in the *envZ*_*R397L*_ was also very high (135 μM) and remained high in the Δ*fur envZ*_*R397L*_ double mutant (105 μM), thus supporting the notion that a higher pool of accessible iron in the *envZ*_*R397L*_ background is responsible for the higher levels of active Fur-Fe^2+^.

**Fig. 3.**
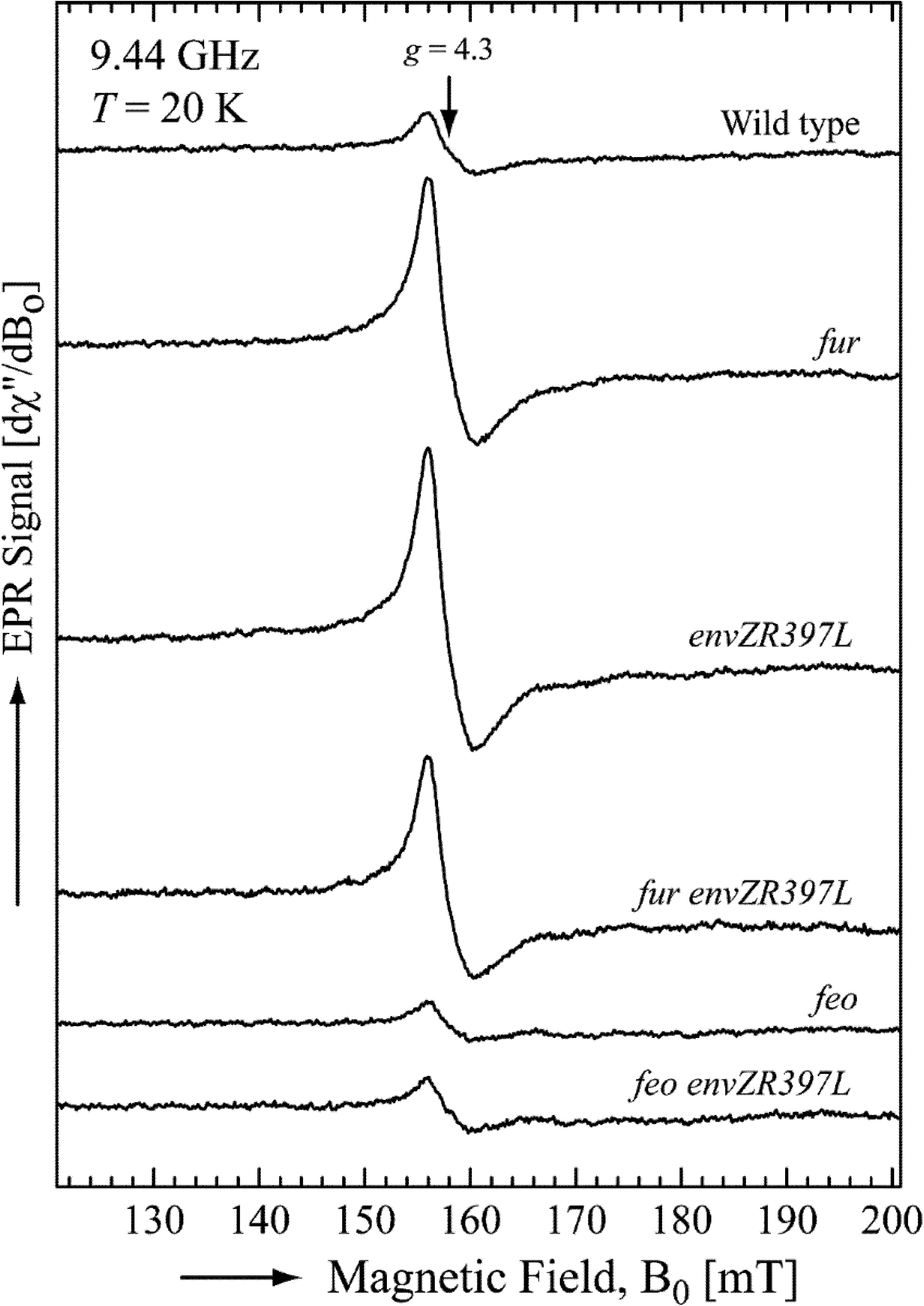
Determination of the intracellular free iron concentration. Shown are the averages of five ferric-chelate EPR scans per strain. All scans were normalized to the final culture OD_600_ used in the measurements. EPR parameters were: microwave power, 10 mW; microwave frequency, 9.44 GHz; center field, 160 mT, sweep width, 80mT; modulation amplitude, 1.25 mT; modulation frequency 100 kHz. Free intracellular iron concentrations, calculated as described in the Experimental procedure section, were: wild type, 32 μM; Δ*fur*, 120 μM; *envZ*_*R397L*_, 135 μM; Δ*fur envZ*_*R397L*_, 105 μM; Δ*feoB*, 20 μM; and Δ*feoB envZ*_*R397L*_, 29 μM. Bacterial strains used are: RAM1292 (wild type), RAM1541 (*envZ*_*R397L*_), RAM2697 (Δ*fur*), RAM2698 (*envZ*_*R397L*_ Δ*fur*), RAM2701 (Δ*feoB*), and RAM2702 (*envZ*_*R397L*_ Δ*feoB*).

Next, we tested whether the FeoB-mediated ferrous transport pathway is responsible for the elevated level of accessible iron in the *envZ*_*R397L*_ mutant. The accessible iron level in the Δ*feoB* mutant was 20 μM or 35% less than the parental *feoB*^+^ strain (Fig. 3), explaining the observed de-regulation of the Fur regulon in the Δ*feoB* mutant (Figs. 1 and 2). Strikingly, without *feoB, envZ*_*R397L*_ failed to increase intracellular iron levels (Fig. 3), thus confirming the involvement of the FeoB-mediated ferrous transport in elevating the intracellular pool of iron, which, in turn, would increase Fur-Fe^2+^ levels and repress expression of *fecA, fepA, fhuA* and *fhuF*. As described below, EnvZ/OmpR play a more direct role in activating *feo* expression to overcome the Fur-mediated downregulation.

### Effects of *envZ*_*R397L*_ on *fepA* and *feo* requires phosphorylated OmpR

Previously it was shown that the pleiotropic effects of the mutant *envZ* allele, *envZ473* with its V241G substitution, is mediated through OmpR (33). In the paper, the authors did not analyze the iron regulon. In this work, we sought to test whether the effect of *envZ*_*R397L*_ on iron regulon requires functional OmpR. We used a missense allele of *ompR* with a D55Y substitution, which confers a null phenotype with respect to *ompC* and *ompF* expression presumably due to the inability of the mutant OmpR to be phosphorylated. The conserved D55 residue of OmpR is the site of phosphorylation (34). The *ompR*_*D55Y*_ allele was isolated in a *fepA*::*lacZ envZ*_*R397L*_ background among Lac^+^ revertants (Misra R., unpublished data). Using a linked Cm^r^ marker, we transduced the *ompR*_*D55Y*_ *envZ*_*R397L*_ mutations into a *feo*::*lacZ* background so that the effects of the mutant *ompR* and *envZ* alleles on *feo* expression can be determined. It is worth noting that although *ompR*/*envZ* are highly linked to the *feo* operon, we were able to construct the above strain since *feo*::*lacZ* is marked by the Km^r^ gene and the mutant *ompR*/*envZ* alleles produce a distinct porin phenotype.

Data presented in Fig. 4 show that *envZ*_*R397L*_ reduced *fepA*::*lacZ* expression about fourfold, whereas *ompR*_*D55Y*_ abolished this effect of *envZ*_*R397L*_. Likewise, the presence of *envZ*_*R397L*_ elevated *feo*::*lacZ* expression fivefold and again *ompR*_*D55Y*_ abolished this increase in *feo* expression. Curiously, *feo*::*lacZ* expression in the *ompR*_*D55Y*_ *envZ*_*R397L*_ background was slightly lower than that seen in the wild type background, suggesting a role for functional OmpR in the expression of the *feo* operon. Together, these data show unambiguously that the negative and positive effects of *envZ*_*R397L*_ on *fepA* and *feo*, respectively, require functional OmpR.

**Fig. 4.**
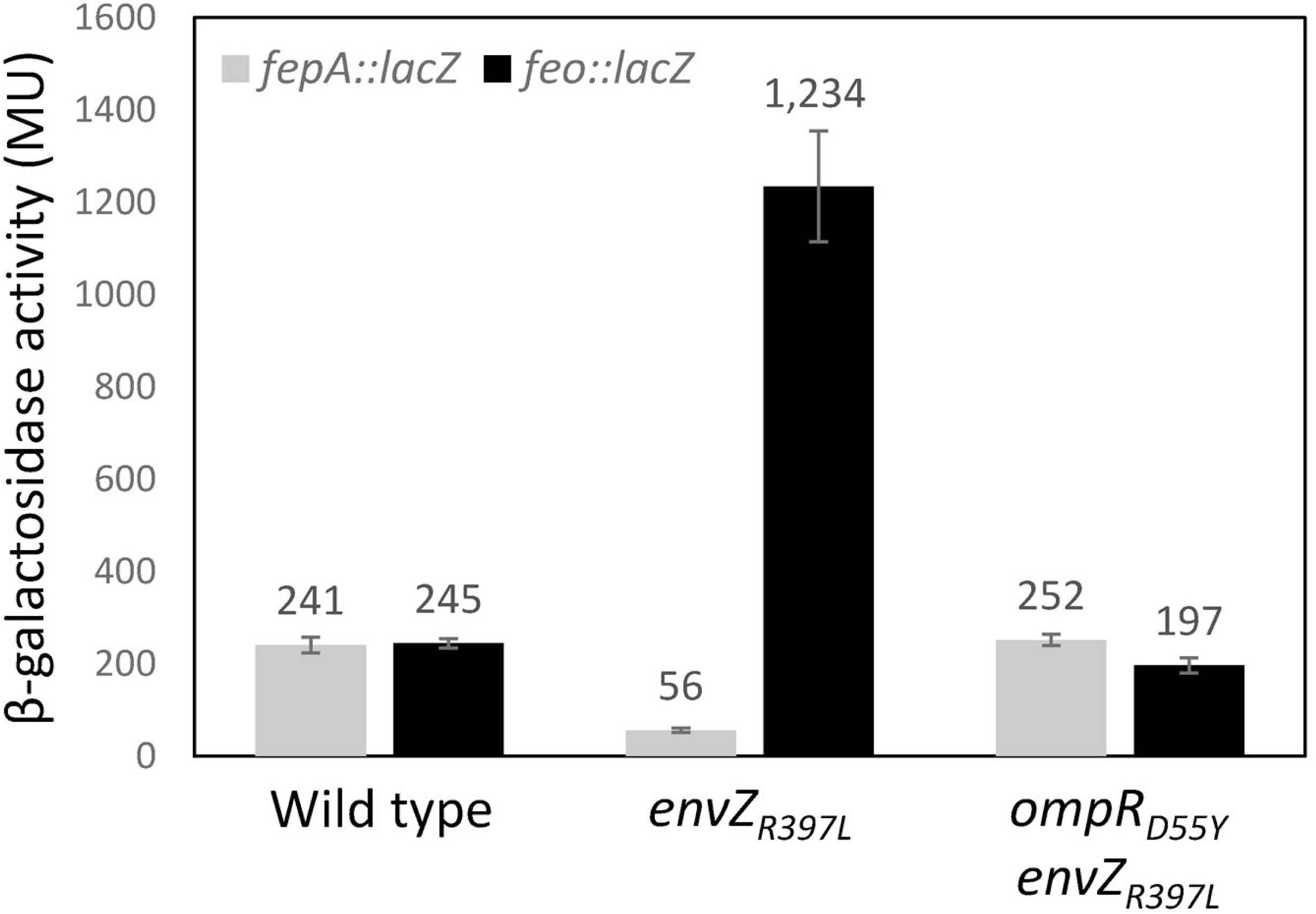
Determination of *fepA*::*lacZ* and *feo*::*lacZ* activities in various genetic backgrounds. The β-galactosidase activities were measured from two independent overnight grown cultures. Error bars represent standard deviation. Bacterial strains used are: RAM2920 (*ompR*^+^ *envZ*^+^ *fepA*::*lacZ*), RAM2921 (*ompR*^+^ *envZ*_*R397L*_ *fepA*::*lacZ*), RAM2922 (*ompR*_*D55Y*_ *envZ*_*R397L*_ *fepA*::*lacZ*), RAM2923 (*ompR*^+^ *envZ*^+^ *feo*::*lacZ*), RAM2924 (*ompR*^+^ *envZ*_*R397L*_ *feo*::*lacZ*), and RAM2925 (*ompR*_*D55Y*_ *envZ*_*R397L*_ *feo*::*lacZ*).

### Direct regulation of feoABC operon by EnvZ/OmpR

The data in Figs. 2 and 4 showed a dramatic increase in the *feo* transcript/transcription levels in the *envZ*_*R397L*_/*ompR*^+^ background. This could be due to the direct regulation of *feo* by OmpR or an effect of an OmpR-controlled factor on the *feo* promoter or *feo* transcript. We took cues from an earlier publication that showed overexpression of RstA, the response regulator of the RstB/RstA TCS, up-regulated *feoB* expression and repressed the Fur regulon in *Salmonella* Typhimurium (35). Electrophoretic mobility shift assays (EMSA) showed direct binding of RstA to the *feo* promoter sequence (Jeon *et al*., 2008). Moreover, the authors identified the “RstA motif” (TACA-N_6_-TACA) upstream of the *S*. Typhimurium *feoA* gene of the *feo* operon (35). Although OmpR recognition sequences are quite degenerate (36; 37), one of the motifs–GTTACANNNN–resembles that of RstA (Fig. 5A). Indeed, both RstA and OmpR regulate some of the same genes by binding to overlapping promoter sequences (38). Our initial assessment detected two potential sequences (−294)-TTATCAtttcaTTAACA-(−278) and (−165)-CCAACAttcgCACACA-(−150) upstream of the *feoA* ATG codon that might contain both RstA and OmpR binding motifs (Fig. 5A).

**Fig. 5.**
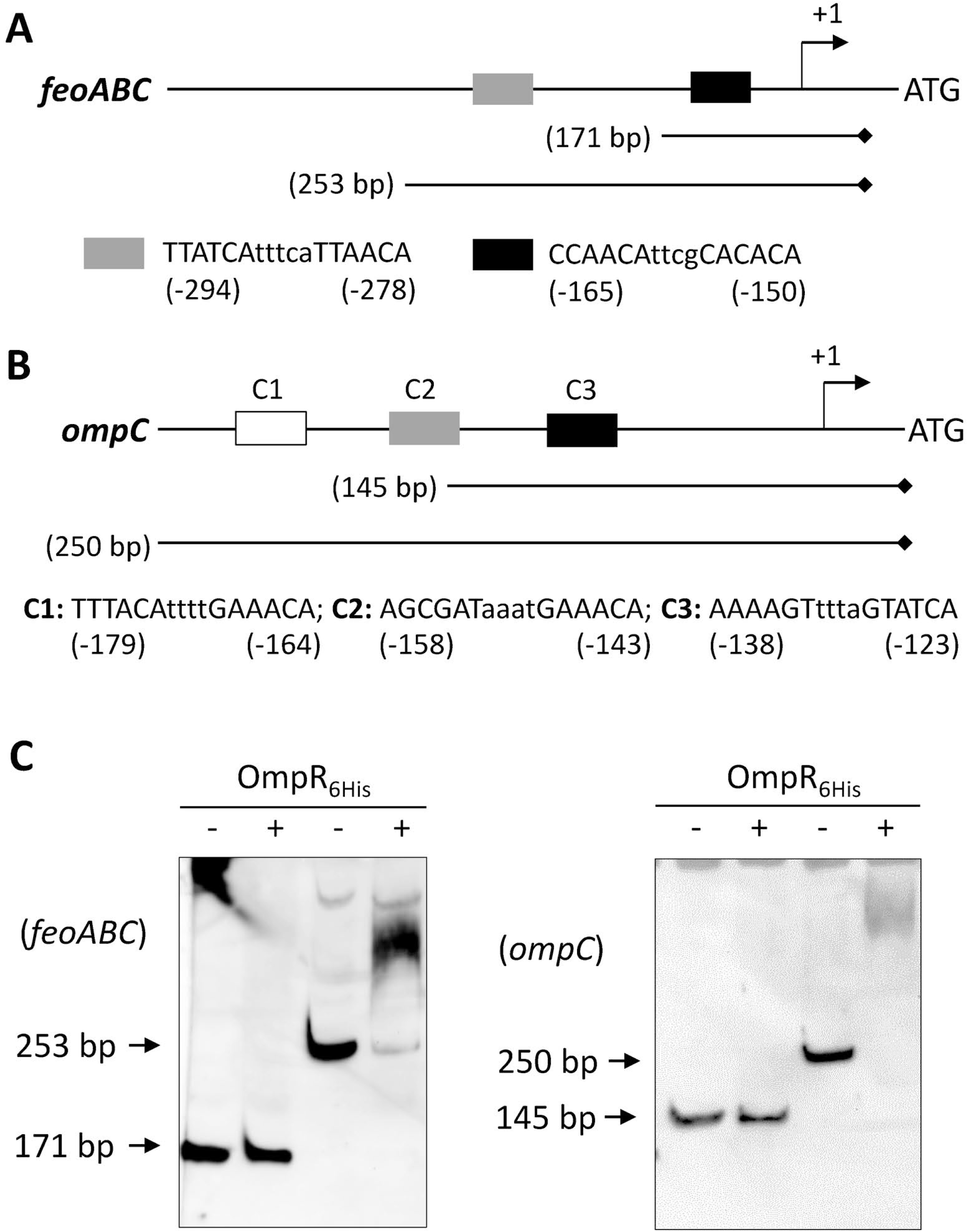
*In vitro* binding of purified OmpR_6His_ to the *feoABC* and *ompC* promoter regions. DNA binding was examined by electrophoretic mobility shift assay (EMSA) using biotin-labeled DNA fragments of various lengths generated by PCR. (**A**) A cartoon showing the regulatory region of the *feoABC* operon (not drawn to scale). Grey and black boxes represent possible OmpR binding sequences. Nucleotide numberings are relative to the *feoA* start codon. Relative positions and lengths of the two DNA fragments used in EMSA are shown. Diamond marks the biotin-labeled end of the DNA probe. (**B**) A cartoon showing the regulatory region of the *ompC* gene (not drawn to scale). Three boxes represent known OmpR binding sites; DNA sequences of all three OmpR binding sites—C1, C2, and C3—are shown. Nucleotide numberings are relative the *ompC* start codon. Relative positions and lengths of the two DNA fragments used in EMSA are shown. Diamond indicates the biotin-labeled end of the DNA probe. (**C**) Polyacrylamide gels showing EMSA results. Plus and minus signs denote the presence and absence of OmpR in the reaction mixture prior to gel electrophoresis. Gels were electroblotted and DNA bands were detected by treating membranes with Stabilized Streptavidin-HRP Conjugate, followed by Luminol/Enhancer and Stable Peroxide. Arrows point to positions of un-shifted DNA fragments.

EMSA was carried out to test whether OmpR can bind directly to the *feo* promoter region. The coding region of *ompR* was cloned into an expression vector, pET24d(+). To aid in protein purification, six consecutive histidine codons were included at the 3’ end of the gene during cloning and the protein was purified to near homogeneity by metal affinity chromatography (Fig. S3). The purified protein was used directly without *in vitro* phosphorylation. Using biotinylated primers, two DNA templates of the *feo* regulatory region, encompassing the predicted OmpR binding motifs, were amplified by PCR (Fig. 5A). As a positive control for OmpR binding, two *ompC* DNA fragments were also included for EMSA (Fig. 5B). No DNA gel shift occurred with the smaller *feoB* DNA fragment containing one of the predicted OmpR binding motifs (Fig. 5C). However, the larger *feo* promoter fragment, containing the upstream predicted OmpR binding motif, displayed shifts after incubation with purified OmpR_6His_ (Fig. 5C). Consistent with these *in vitro* data, we found that overexpression of OmpR_His_ from a pBAD24 replicon increased *feo*::*lacZ* expression twofold (from 140±8 Miller units in the pBAD24 vector containing strain to 296±25 Miller units in the strain containing pBAD24-*ompR*_*His*_). OmpR bound to the *ompC* promoter fragment containing the high-affinity OmpR-binding motif C1 (39), but not with the one containing the partial C2 and the entire C3 motif (Fig. 5C). Incidentally, only the *ompC* fragment, containing all three OmpR motifs, expressed the promoter-less *lacZ* gene in an OmpR-dependent manner (Fig. S4), thus corroborating the EMSA data. Together these data indicated that OmpR positively regulates *feo* expression by directly binding to the *feo* promoter region.

### Role of porins in iron homeostasis

The data in Figs. 1-4 revealed a possible mechanism by which a pleiotropic allele of *envZ* downregulates the ferric transport systems by employing the OmpC/FeoB-mediated ferrous transport pathway. While these data implicated EnvZ/OmpR and OmpC in iron transport, the use of a pleiotropic *envZ* allele may have created an unnatural genetic environment in which EnvZ/OmpR and porins become involved in iron homeostasis. To eliminate this possibility, we determined the roles EnvZ/OmpR and porins in iron transport using the null alleles of *ompR* and the porin genes. Before testing their roles, we disabled the high-affinity ferric transport system, since porins likely mediate iron transport by simple diffusion of ferrous or small iron-chelated compounds and this passive activity of porins will likely be masked by the high-affinity iron transport system. In *E. coli*, the high-affinity iron transport principally involves a ferric chelator, enterobactin, and TonB that interacts with the outer membrane iron receptors for the release of chelator-Fe^3+^ complexes bound to the receptor. Consequently, we disabled the ferric iron transport by deleting *aroB, tonB* or both. The *aroB* gene encodes 3-dehydroquinate synthase, which is required for the second step of the chorismate pathway in the synthesis of enterobactin, aromatic amino acids and other important compounds (40).

We first determined the iron dependency of wild type, Δ*aroB*, Δ*tonB*, and Δ*aroB* Δ*tonB* strains by growing them on LBA, LBA supplemented with 40 μM FeCl_3_ and LBA containing 200 μM of 2,2’-dipyridyl (DP), a synthetic iron chelator (Fig. 6A-C). Bacterial growth in the absence of *aroB* was unaffected on LBA+FeCl_3_ or LBA (Fig. 6A, B). However, significant growth impairment of the Δ*aroB* strain occurred on LBA+DP plates (Fig. 6C), reflecting the loss of a major, enterobactin-mediated iron transport system. In contrast to Δ*aroB*, the deletion of *tonB* impaired bacterial growth even on LBA (Fig. 6B), which contains around 6 μM of iron, and completely prevented growth on LBA+DP medium (Fig. 6C). The Δ*tonB* strain grew like WT on LBA+FeCl_3_, showing that the growth impairment of this strain on LBA was due to low accessibility to iron. Interestingly, growth of the Δ*aroB* Δ*tonB* double mutant improved slightly on LBA compared to the Δ*tonB* strain (Fig. 6B), but ceased again on LBA+DP (Fig. 6C). An improvement in growth of the double mutant compared the Δ*tonB* strain on LBA may be due to the absence of extracellular enterobactin-Fe^3+^ complexes, which, when allowed to accumulate outside the Δ*tonB* cells, would sequester iron from the medium and further exacerbate growth defects (41). Because of the greater growth dependence of the Δ*tonB* and Δ*tonB* Δ*aroB* strains on external iron sources than the Δ*aroB* strain, we selected the former two genetic backgrounds to examine the effects of EnvZ/OmpR and porins in iron transport. It is worth noting that we did not determine bacterial growth rates by monitoring growth of liquid cultures because the Δ*tonB* strain frequently reverts without supplemented iron, and these faster growing revertants takeover the population to artificially display better than expected growth.

**Fig. 6.**
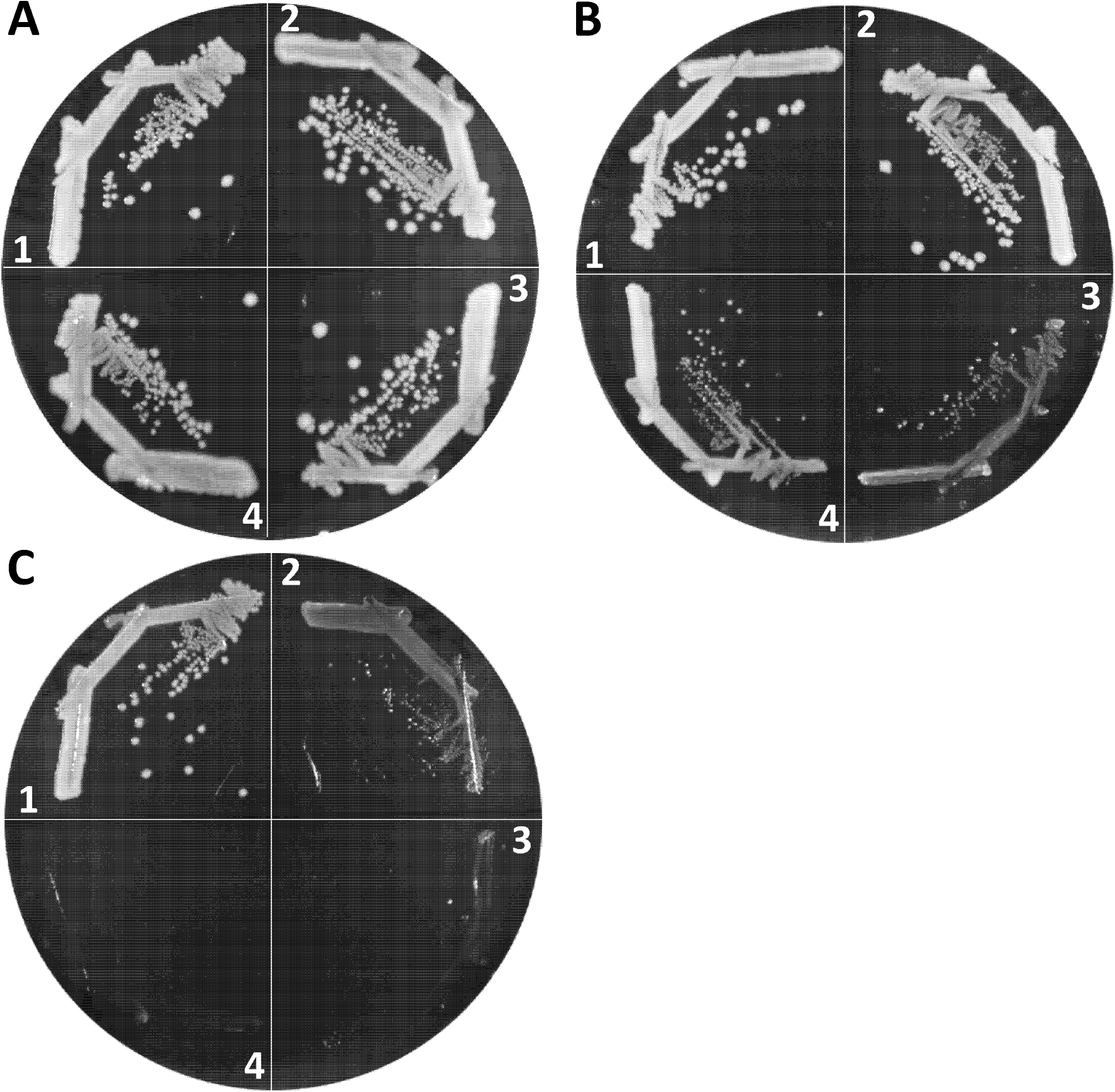
Effects of Δ*tonB* and Δ*aroB* mutations on bacterial growth under iron replete and deplete conditions. Bacterial growth on LBA + 40 μM FeCl_3_ (**A**), LBA (**B**), and LBA+200 μM 2,2’-dipyridyl (**C**) was recorded after incubating petri plates at 37°C for 24 h. Bacterial strains used are: 1, RAM1292 (wild type); 2, RAM2553 (Δ*aroB*); 3, RAM2572 (Δ*tonB*) and 4, RAM2574 (Δ*aroB* Δ*tonB*).

We employed two different null *ompR* alleles, *ompR101* and Δ*ompR*::Km^r^, both of which produce the OmpC^-^ OmpF^-^ phenotype. The *ompR101* allele was transduced into a Δ*tonB* background using a linked Tc^r^ marker, *malPQ*::Tn*10*, while Δ*ompR*::Km^r^ was transduced directly using the Km^r^ gene that replaced the deleted *ompR* gene. Although both *ompR* alleles could be transduced in the Δ*tonB* strain when transductants were selected on LBA+FeCl_3_ containing appropriate antibiotics, the resulting null *ompR* Δ*tonB* transductants grew poorly compared to the *ompR*^+^ Δ*tonB* strain (Fig. 7A; sectors 4 and 5). In contrast, *ompR101* and Δ*ompR*::Km^r^ severely compromised growth of the Δ*tonB* strain on LBA not supplemented with FeCl_3_ (Fig. 7B; sectors 4 and 5). Similar to the Δ*tonB ompR101* strain, we were able to construct the Δ*aroB* Δ*tonB ompR101* strain on LBA+FeCl_3_ medium, where it grew poorly (Fig. 7A; sector 8) but not as poorly as on LBA without FeCl_3_, where the strain failed to form single colonies (Fig. 7B; sector 8). These observations pointed to a critical role for the EnvZ/OmpR TCS in iron transport in the absence of the high-affinity iron transport system.

**Fig. 7.**
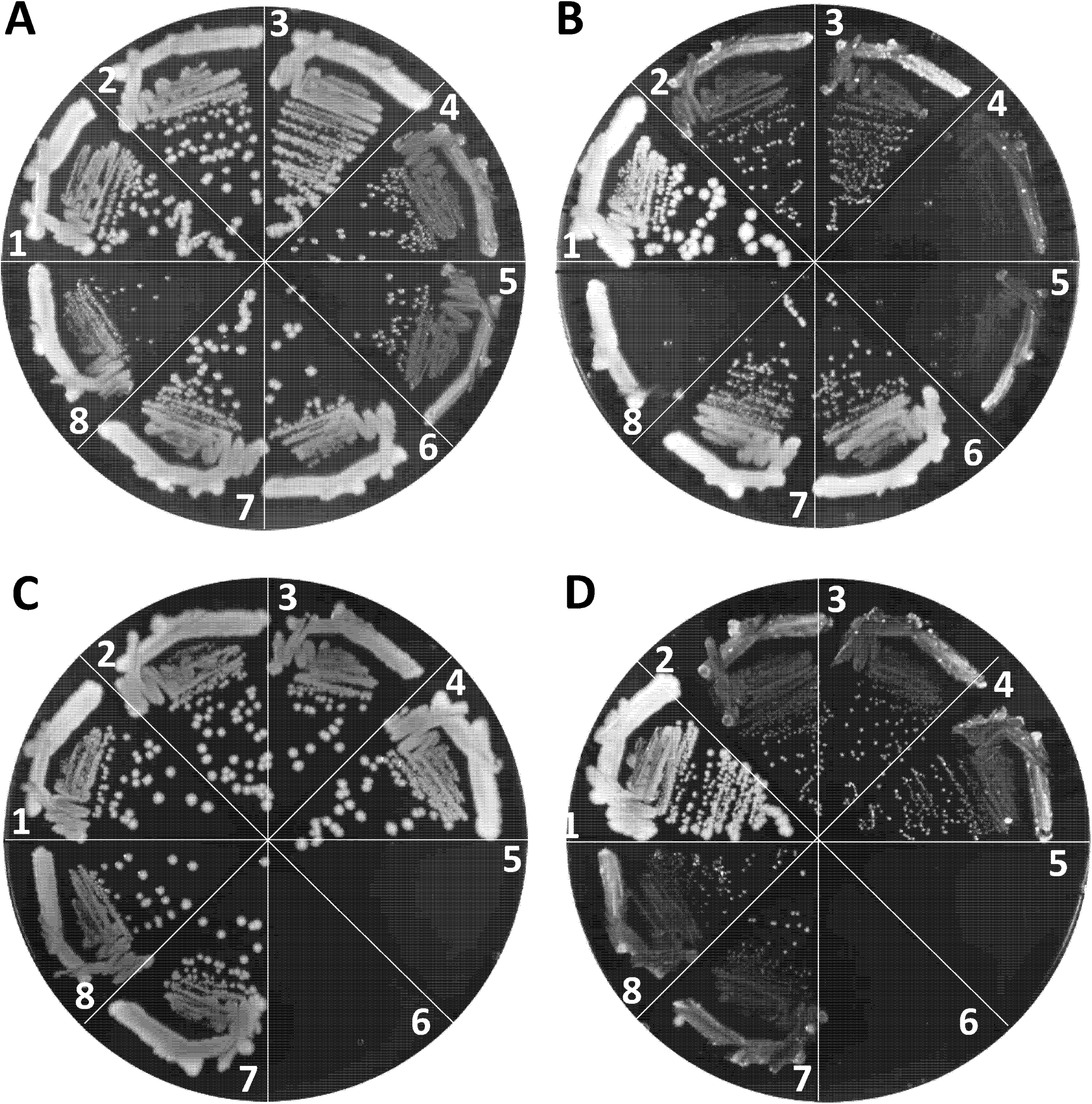
Effects of *ompR* and the porin gene mutations on the growth of Δ*tonB* or Δ*tonB* Δ*aroB* mutants. Bacterial growth was monitored on LBA + 40 μM FeCl_3_ (**A** and **C**) and LBA (**B** and **D**) after incubating petri plates at 37°C for 24 h. The relevant genotypes of strains used in **A** and **B** are: 1, RAM1292 (wild type); 2, RAM2572 (Δ*tonB*); 3, RAM2765 (Δ*tonB malPQ*::Tn*10*); 4, RAM2766 (Δ*tonB malPQ*::Tn*10 ompR101*); 5, RAM2767 (Δ*tonB* Δ*ompR*::Km^r^); 6. RAM2574 (Δ*tonB* Δ*aroB*::Km^r^); 7, RAM2771 (Δ*tonB* Δ*aroB*::Km^r^ *malPQ*::Tn*10*); and 8, RAM2772 (Δ*tonB* Δ*aroB*::Km^r^ *malPQ*::Tn*10 ompR101*). The relevant genotypes of strains used in **C** and **D** are: 1, RAM1292 (wild type); 2, RAM2572 (Δ*tonB*); 3, RAM2769 (Δ*tonB* Δ*ompC*::Cm^r^); 4, RAM2768 (Δ*tonB* Δ*ompF*::Km^r^); 5 and 6, no bacteria; 7, RAM2792 (Δ*tonB* Δ*ompC*::Cm^r^ Δ*ompF*::Km^r^/p*ompC*); and 8, RAM2790 (Δ*tonB* Δ*ompF*::Km^r^ Δ*ompC*::Cm^r^/p*ompF*). p*ompF* and p*ompC* are pTrc99A plasmid clones expressing *ompF* and *ompC*, respectively. Expression of these plasmid-coded genes did not require induction by an inducer.

Although the porin genes are the main targets of the EnvZ/OmpR regulatory system, transcription of other genes is also affected either directly or indirectly in the *ompR* null mutant (26). Therefore, to establish unambiguously the importance of porins in iron transport we attempted to delete the porin genes in a background devoid of the high-affinity transport system. In the Δ*tonB* background, the deletion of *ompC* or *ompF* individually did not significantly influence growth on LBA (Fig. 7C, D; compare sectors 3 and 4 with 2). Strikingly, however, we failed to delete *ompC* and *ompF* simultaneously, via P1 transduction of Δ*ompF*::Km^r^ and Δ*ompC*::Cm^r^ alleles, in the Δ*tonB* background even when transductants were selected on LBA+FeCl_3_ plates carrying appropriate antibiotics. In contrast, when the Δ*tonB* Δ*ompC* or Δ*tonB* Δ*ompF* double mutant was first complemented with a plasmid expressing one of the porin genes, the un-complemented porin gene from the chromosome could be readily deleted by P1 transduction. The plasmid-complemented triple mutants displayed growth behavior similar to the un-complemented Δ*tonB* Δ*ompC* and Δ*tonB* Δ*ompF* double mutants on LBA+FeCl_3_ or LBA (Fig. 7C, D; compare sectors 3 and 4 with 7 and 8). It is worth noting that the Δ*tonB* Δ*ompC* and Δ*tonB* Δ*ompF* strains were not defective in P1 transduction, since drug resistant markers not associated with the porin genes or their regulators could be transduced readily into these strains. Moreover, unlike the Δ*tonB* strain, in the wild type and Δ*aroB* backgrounds the *ompC* and *ompF* genes could be deleted simultaneously without causing iron dependency or a significant growth defect (Fig. S5). These data indicated, for the first time, that the OmpC and OmpF porins play a critical role in iron intake when the high-affinity iron transport system is blocked.

## DISCUSSION

Although the EnvZ/OmpR TCS is classically associated with the regulation of the OmpC and OmpF porins in response to medium osmolarity (23), recent transcriptomics and chromatin immunoprecipitation analyses showed that it is a global regulatory system (26, 37, 42). Indeed, missense alleles of *envZ*, called *perA* and *tpo*, isolated over three decades ago, were shown to also influence non-porin regulons, including *pho* and *mal* (43, 44, 45). A separate study revealed that *perA* lowered the expression of three iron-regulated proteins without an apparent reduction in the rate of enterobactin secretion (27). This led the authors to suggest that the effect of the *perA* (*envZ*) allele on the expression of iron-regulated proteins is most likely post-transcriptional (27).

In this study, we sought to resolve the mechanism by which the activated EnvZ/OmpR TCS reduces expression of genes involved in iron homeostasis and determine the role of porins in iron acquisition. We used the *envZ*_*R397L*_ allele, which is phenotypically similar to the pleotropic *perA* and *tpo* alleles of *envZ*, i.e., in the *envZ*_*R397L*_ background, OmpC levels go up, while those of OmpF and LamB go down dramatically (29). The RT-qPCR (this work) and the whole genome microarray data (30; Table S1) showed that in the presence of *envZ*_*R397L*_, transcript levels of several Fur-controlled genes, including *fecA, fepA, fhuA* and *fhuF*, went down significantly. In the case of *fhuA* and *fhuF*, the effects of *envZ*_*R397L*_ required Fur, while expression of *fecA* and *fepA* was still reduced by *envZ*_*R397L*_ in the absence of Fur. These observations indicated the involvement of at least two different mechanisms by which *envZ*_*R397L*_ affected iron regulon. In support of the Fur-dependent mechanism, the whole-cell EPR data confirmed the presence of significantly elevated levels of accessible iron in the *envZ*_*R397L*_ strain. Several observations supported the hypothesis that in the *envZ*_*R397L*_ mutant, FeoB and OmpC are responsible for increased intracellular Fur-Fe^2+^ level (Fig. 8). First, unlike the expression of genes involved in ferric iron transport or metabolism, expression of the *feoAB* genes involved in ferrous iron transport went up dramatically in the *envZ*_*R397L*_ background. This increase in the expression of the ferrous iron transport system had an adverse effect on the ferric iron transport system, since the absence of FeoB, the ferrous permease, abolishes or significantly reduces the negative effects of *envZ*_*R397L*_ on ferric transport/metabolic genes. Second, like FeoB, the absence of OmpC (*envZ*_*R397L*_ already severely represses *ompF* expression; 29) largely negated the inhibitory effects of *envZ*_*R397L*_ on ferric transport/metabolic genes. Because the single deletion of *feoB* or *ompC* and the simultaneous deletion of *feoB* and *ompC* reversed the effects of *envZ*_*R397L*_ on *fecA, fepA, fhuA* and *fhuF* to the same extent, it indicated that FeoB and OmpC must act in the same pathway to transport ferrous iron into the cell and elevate Fur-Fe^2+^ levels. Third, the absence of FeoB or OmpC in an EnvZ^+^ background caused de-repression of six Fur-controlled genes, indicating that the ferrous iron transport pathway is active under our experimental conditions and that *envZ*_*R397L*_ enhances this pathway to achieve its inhibitory effects on the ferric transport system. Lastly, we provided direct evidence of excessive iron inside the *envZ*_*R397L*_ mutant by whole-cell EPR spectroscopy measurements, which showed that, as in the Δ*fur* mutant, the level of accessible iron in the *envZ* mutant rose fourfold over that present in the parental strain. Moreover, this increase in the intracellular free pool of iron in the *envZ*_*R397L*_ mutant was dependent on FeoB. From these observations, we conclude that the upregulation of the OmpC-FeoB ferrous iron transport pathway by *envZ*_*R397L*_ elevates the intracellular Fur-Fe^2+^ level, which, in turn, represses the expression of iron-regulated genes (Fig. 8). These effects of *envZ*_*R397L*_ required functional OmpR since the presence of *ompR*_*D55Y*_, which confers a null phenotype, neutralized all *envZ*_*R397L*_ phenotypes.

**Fig. 8.**
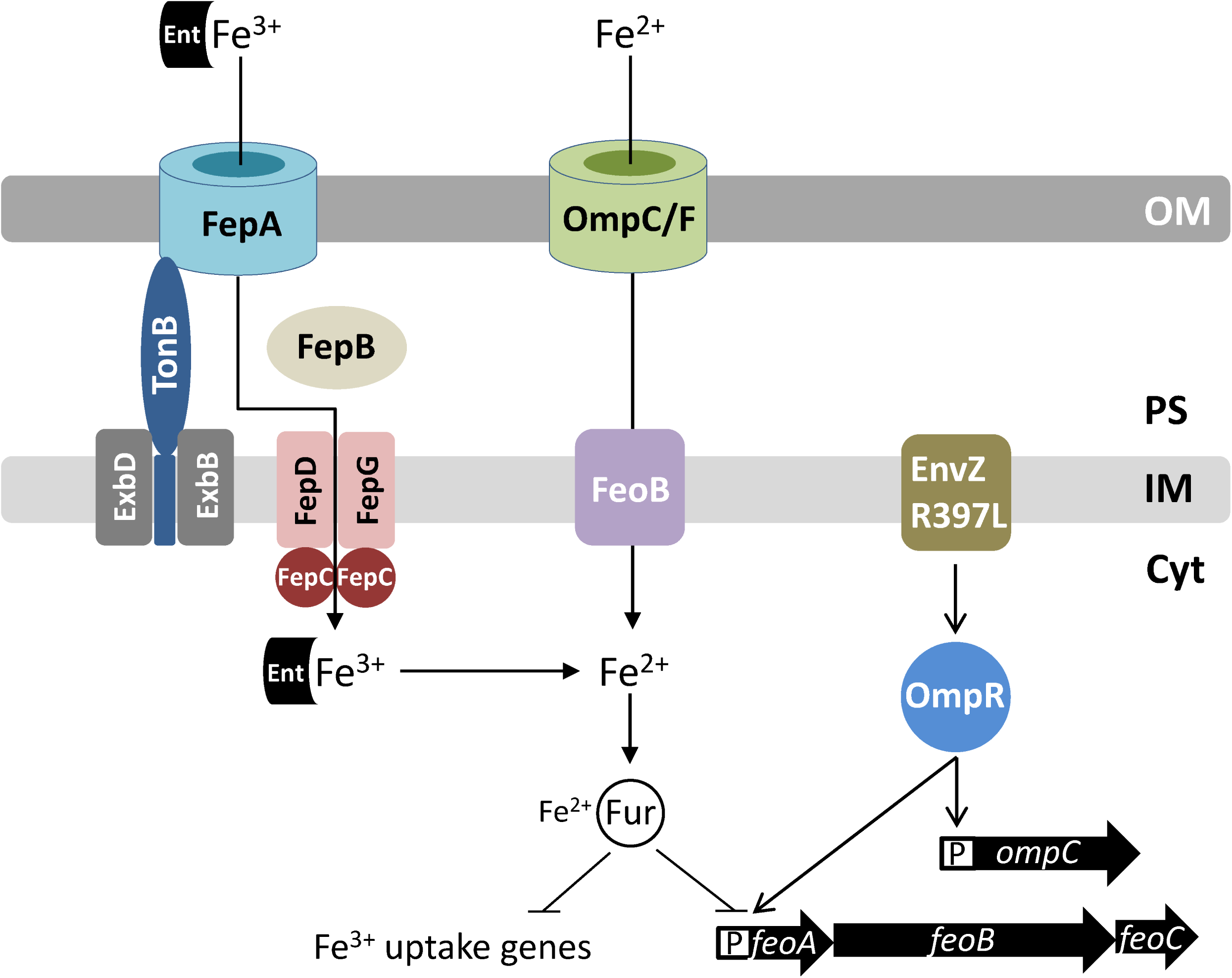
A cartoon showing regulation of the ferric (Fe^3+^) and ferrous (Fe^2+^) uptake systems in *E. coli*. Fur-Fe^2+^ is the master regulator of transcription of genes involved in iron metabolism. Under aerobic growth conditions, where Fe^3+^ is the major source of iron, *E. coli* secretes enterobactin (Ent) in the medium to chelate Fe^3+^. The Fe^3+^-chelate complex is transported back into the cell through the outer membrane receptor protein, FepA. The TonB-ExbB-ExbD complex of the inner membrane facilitate FepA channel opening. In the periplasm, FepB interacts with the Fe^3+^-chelate and delivers it to the FepDGC complex for transport into the cytoplasm. Under micro-aerobic or anaerobic growth conditions, Fe^2+^ is the main source of iron. It is brought into the cell via porins OmpC and OmpF and FeoB. The EnvZ/OmpR two-component system, classically known for regulating the expression of the *ompC* and *ompF* porin genes, also induces *feo* expression when hyper-activated due to a specific mutation in *envZ* (*envZ*_*R397L*_). This positive effect of EnvZ_R397L_/OmpR on *feo* expression can overcome the negative effect of Fur-Fe^2+^ on *feo* expression, thus tipping the balance in favor of ferrous over ferric transport. Porins and EnvZ/OmpR play a crucial role in iron acquisition in a TonB-deficient background that lacks functional ferric transport systems. Abbreviation: OM, outer membrane; PS, periplasm; IM, inner membrane; and Cyt, cytoplasm; and P, promoter.

Whereas *envZ*_*R397L*_-mediated reduction in *fhuA* and *fhuF* transcript levels required Fur, the effects of *envZ*_*R397L*_ on *fepA* and *fecA* transcripts did not. This suggested the existence of another regulatory mechanism responsible for the *envZ*_*R397L*_-mediated downregulation of *fepA* and *fecA* that did not involve Fur. Previous studies showed that the plasmid-mediated overexpression of OmrA and OmrB small RNAs, whose expression is under the EnvZ/OmpR control, can downregulate *fepA* and *fecA* transcript levels (31, 46). We have shown that *envZ*_*R397L*_ increases OmrA expression almost tenfold (29). This increase in OmrA expression could contribute to the downregulation of *fepA* and *fecA*. However, the fact that deleting *ompC* or *feoB* in a Fur^+^ background abolishes *envZ*_*R397L*_-mediated downregulation of the ferric iron transport genes suggests that *envZ*_*R397L*_-mediated increase in OmrA and OmrB levels contributes little, if any, to *fepA* and *fecA* repression. Consistent with this notion, we found that the deletion of Δ*omrA* and Δ*omrB* failed to reverse the negative effect of *envZ*_*R397L*_ on *fepA* or *fecA* (Fig. S2). We conclude that a mechanism independent of Fur and OmrA and OmrB must also exist for the *envZ*_*R397L*_-mediated downregulation of *fepA* and *fecA*. A direct role of EnvZ/OmpR has not been ruled out.

As stated above, unlike the ferric transport genes, expression of the ferrous transport genes *feoAB*, which are also under the control of Fur, went up in the *envZ*_*R397L*_ background. At first glance, this appears inconsistent with the notion that an increase in the Fur-Fe^2+^ level by *envZ*_*R397L*_ should also decrease *feoAB* expression. Our data suggest that *envZ*_*R397L*_ overcomes the repressive effect of Fur-Fe^2+^ on *feoAB* by activating their expression. Moreover, because *feoAB* expression in the *envZ*_*R397L*_ background increases dramatically without Fur, it shows that Fur-Fe^2+^ does repress *feoAB* in the *envZ*_*R397L*_ background, but the positive effect of *envZ*_*R397L*_ on *feoAB* expression overwhelms the negative effect of Fur on these genes. The EMSA data showed that purified OmpR binds to the *feo* regulatory region containing a putative OmpR binding site, indicating that OmpR directly activates *feoAB* expression. Interestingly, the whole genome microarray data from an Δ*ompR* Δ*envZ* (porin-minus) strain showed that *feoB* transcript levels decreased twofold, whereas those of *fepA* and *fecA* went up twofold (26). These observations are consistent with our proposal that the EnvZ/OmpR TCS directly stimulates the FeoB-OmpC pathway to increase intracellular Fe^2+^ levels and thus active Fur-Fe^2+^ complexes, which then downregulate the expression of the ferric iron transport genes.

The proposed role of EnvZ/OmpR in iron homeostasis is similar to that suggested for RstA in *S*. Typhimurium (35). The authors found that overexpression of RstA increased *feoAB* expression and repressed *fhuA* and *fhuF* expression. A RstA binding site was identified in the *feo* promoter and the EMSA data confirmed that RstA bound there (35). The RstA binding motif ‘TACAtntngtTACA’ resembles that of OmpR’s ‘GTTACAnnnnGTTACA’ and not surprisingly, both proteins regulate overlapping genes by binding to the similar sequences (38). Our EMSA data showed that OmpR binds to the *feo* promoter region. Specifically, it binds to a DNA fragment containing the sequence ‘ttATCAtttcattAACA’ located 278 bp upstream of the start codon of *feoA*. The OmpR binding studies carried out here involved the purified protein not modified by *in vitro* phosphorylation. Therefore, it is possible that stronger binding and/or additional binding sites may be discovered with phosphorylated OmpR. It is worth noting that in previous EMSA assays, unphosphorylated RstA from *S*. Typhimurium and *E. coli* was shown to bind to their target promoter sequences (35, 47). Further work will be required to identify the exact binding sequences and to determine whether OmpR and RstA bind to the same, overlapping or distinct regulatory sequences of the *feo* operon.

Our work also revealed, for the first time, the essential role of OmpC and OmpF porins in iron acquisition when the TonB-dependent ferric transport pathways are inoperative. The absence of OmpC or OmpF produced no growth defects in the Δ*tonB* background on LBA supplemented with iron, but the construction of a triple knockout mutant (Δ*tonB* Δ*ompC* Δ*ompF*) required expression of at least one of the porin genes from a plasmid replicon. Interestingly, unlike the porin-devoid triple knockout mutant, we were able to construct Δ*tonB* Δ*ompR* and Δ*tonB* Δ*aroB* Δ*ompR* mutants albeit, only on LBA supplemented with iron. In the Δ*ompR* background, *ompC* and *ompF* porin expression is extremely low but presumably not zero, which is the case in the Δ*tonB* Δ*ompC* Δ*ompF* mutant. We think that this extremely low porin expression permits the construction of the Δ*tonB* Δ*ompR* strain, which can form single, albeit very small colonies, but only on LBA supplemented with iron. Whereas the diffusion of ferrous iron across the outer membrane occurs via OmpC or OmpF channels, at least three proteins— FeoB, MntH and ZupT—can transport ferrous iron across the inner membrane of *E. coli* cells (48). Consistent with this, a Δ*tonB* Δ*feoB* double mutant is viable and grows like the Δ*tonB* mutant (data not shown).

While the essential role of porins in iron acquisition becomes apparent without the TonB-dependent, high-affinity ferric iron transport systems, their de-repression without OmpC or FeoB indicate that the porin-mediated iron transport is active even in the presence of the TonB-dependent high-affinity iron transport systems. The importance of the porin-FeoB pathway for bacterial growth should further increase as *E. coli* cells enter microaerobic or anaerobic environments where the ferrous species predominates. The involvement of porin and FeoB in iron-dependent growth and/or virulence has been reported for several bacteria, including *E. coli* (9), *S*. Typhimurium (10; 11), *Helicobacter pylori* (12), *Vibrio cholerae* (49) and *Mycobacterium smegmatis* (50). Interestingly, *M. smegmatis* porins increase ferric citrate uptake (50). Similarly, a study reported liganded iron uptake via the OprF porin in *Pseudomonas aeruginosa* (51). There are no definitive reports in *E. coli* showing the involvement of porins in liganded iron transport, even for ferric citrate whose size is below the diffusion limits of the porins (52). Regardless of these ambiguities, published reports and the work carried out here highlight the importance of the porin/FeoB-mediated iron transport pathways in iron homeostasis.

## MATERIALS AND METHODS

### Bacterial strains, media and chemicals

*Escherichia coli* K-12 strains used in this study were constructed from MC4100 (53) and are listed in Table 1. Luria broth (LB) was prepared using LB Broth EZMix(tm) Powder (Lennox). LB agar (LBA) medium contained LB plus 1.5% agar (Becton Dickenson). ONPG (2-ortho-nitrophenyl-β-D-galactopyranoside) was purchased from ACROS. Diethylenetriaminepentaacetic acid (DTPA) and desferrioxamine were obtained from Sigma-Aldrich. All other chemicals were of analytical grade. The growth medium was supplemented with ampicillin (50 μg/ml), chloramphenicol (12.5 μg/ml), kanamycin (25 μg/ml) or tetracycline (10 μg/ml) when necessary. To induce plasmid-borne gene expression, L-arabinose (0.2%) or isopropyl β-D-1-thiogalactopyranoside (IPTG; 0.4 mM) was added to the medium.

**Table 1.** Bacterial strains used in this study.

| Strain | Relevant genotype | Source |
| --- | --- | --- |
| RAM1292 | MC4100 ($\Delta[argF-lac]169 \lambda^- e14^- flhD5301 \Delta[fruK-yeiR]725 relA1 rpsL150 (str^R) rbsR22 \Delta[fimB-fimE]632 deoC1) \Delta ara714$ | 61 |
| RAM1541 | RAM1292 *envZ*<sub>R397L</sub> | 29 |
| RAM2697 | RAM1292 $\Delta fur::scar$ | This study |
| RAM2698 | RAM1292 *envZ*<sub>R397L</sub> $\Delta fur::scar$ | This study |
| RAM2699 | RAM1292 $\Delta ompC::scar$ | This study |
| RAM2700 | RAM1292 *envZ*<sub>R397L</sub> $\Delta ompC::scar$ | This study |
| RAM2701 | RAM1292 $\Delta feoB::scar$ | This study |
| RAM2702 | RAM1292 *envZ*<sub>R397L</sub> $\Delta feoB::scar$ | This study |
| RAM2703 | RAM1292 *envZ*<sub>R397L</sub> $\Delta feoB::scar \Delta ompC::scar$ | This study |
| RAM2704 | RAM1292 $\Delta ompR::scar$ | This study |
| RAM2705 | RAM1292 $\Delta ompR::scar$ pBAD24-*ompR*<sub>6His</sub> | This study |
| RAM2707 | RAM1292 $\Delta ompR::scar$ pBAD24 | This study |
| RAM2708 | RAM1292 pBAD24 | This study |
| RAM2709 | RAM1292 pBAD24-*ompR*<sub>6His</sub> | This study |
| RAM2711 | RAM1292 $\Delta feoA::lacZ-Km^r$ | This study |
| RAM2712 | RAM1292 $\Delta feoA::lacZ-Km^r envZ_{R397L}$ | This study |
| RAM2713 | RAM1292 $\Delta feoA::lacZ-Km^r \Delta fur::scar$ | This study |
| RAM2714 | RAM1292 $\Delta feoA::lacZ-Km^r \Delta fur::scar envZ_{R397L}$ | This study |
| RAM2715 | BL21(DE3) pET24d-*ompR*<sub>6His</sub> | This study |
| RAM2469 | RAM1292 $\Delta fepA::lacZ-Km^r$ | This study |
| RAM2470 | RAM1292 $\Delta fepA::lacZ-Km^r envZ_{R397L}$ | This study |
| RAM2471 | RAM1292 $\Delta fepA::lacZ-Km^r \Delta fur$ | This study |
| RAM2472 | RAM1292 $\Delta fepA::lacZ-Km^r \Delta fur envZ_{R397L}$ | This study |
| RAM2473 | RAM2469 $\Delta ompC::Cm^r$ | This study |
| RAM2474 | RAM2469 $\Delta feoB::Cm^r$ | This study |
| RAM2475 | RAM2470 $\Delta ompC::Cm^r$ | This study |
| RAM2476 | RAM2470 $\Delta feoB::Cm^r$ | This study |
| RAM2477 | RAM2469 $\Delta ompF::Cm^r$ | This study |
| RAM2478 | RAM2470 $\Delta ompF::Cm^r$ | This study |
| RAM2505 | RAM1292 $\Delta tonB::Km^r$ | This study |
| RAM2553 | RAM1292 $\Delta aroB::Km^r$ | This study |
| RAM2572 | RAM2505 $\Delta tonB::scar$ (via pCP20) | This study |
| RAM2574 | RAM2572 $\Delta aroB::Km^r$ | This study |
| RAM2623 | RAM2553 $\Delta aroB::scar$ | This study |
| RAM2625 | RAM2574 $\Delta aroB::scar$ (via pCP20) | This study |
| RAM2765 | RAM2572 *malPQ::Tn10-ompR*<sup>+</sup> | This study |
| RAM2766 | RAM2572 *malPQ::Tn10-ompR101* | This study |
| RAM2767 | RAM2572 $\Delta ompR::Km^r$ | This study |
| RAM2768 | RAM2572 $\Delta ompF::Km^r$ | This study |
| RAM2769 | RAM2572 $\Delta ompC::Cm^r$ | This study |
| RAM2771 | RAM2574 *malPQ::Tn10-ompR*<sup>+</sup> | This study |
| RAM2772 | RAM2574 *malPQ::Tn10-ompR101* | This study |
| RAM2790 | RAM2768 (pTrc99A-*ompF*) | This study |
| RAM2792 | RAM2769 (pTrc99A-*ompC*) | This study |
| RAM2795 | RAM2790 $\Delta ompC::Cm^r$ | This study |
| RAM2796 | RAM2792 $\Delta ompF::Km^r$ | This study |
| RAM2920 | RAM2469 $Cm^r$ linked to $envZ^+$ | This study |
| RAM2921 | RAM2470 $Cm^r$ linked to $envZ_{R397L}$ | This study |
| RAM2922 | RAM2799 $Cm^r$ linked to $ompR_{D55Y} envZ_{R397L}$ | This study |
| RAM2924 | RAM2711 $Cm^r$ linked to $envZ^+$ | This study |
| RAM2925 | RAM2711 $Cm^r$ linked to $envZ_{R397L}$ | This study |
| RAM2926 | RAM2711 $Cm^r$ linked to $ompR_{D55Y} envZ_{R397L}$ | This study |
| RAM2928 | RAM2711 pBAD24 ($Ap^r$) | This study |
| RAM2932 | RAM2711 pBAD24-$ompR_{His}$ | This study |

### Genetic and DNA methods

Standard bacterial genetic methods, including P1 transduction and plasmid transformation, were carried out as described by Silhavy *et al*. (54). To clone the *ompR* and *rstA* genes into pBAD24 (55) and pET24 d(+) (Novagen), DNA corresponding to their open reading frames (ORFs) were amplified by PCR using primers that carried appropriate restriction enzyme sites for cloning. The reverse primers used for cloning into pBAD24 additionally contained nucleotides encoding six consecutive histidine codons. (Primer sequences are available upon request.) Deletion of the *fepA, feoA, feoB* and *feoAB* genes from their chromosomal locations and subsequent scaring of the antibiotic-resistant marker at the deletion sites were done using the λ-red-mediated gene recombination method (56). Deletions were confirmed by PCR and DNA sequence analyses. In some instances, promoter-less *lacZY* were recombined at the deletion scar site by the method of Ellermeier *et al*. (57).

### RNA isolation, real-time quantitative PCR, and microarray analyses

Total RNA was extracted from 5 ml cells grown to log phase (OD_600_ ∼0.6) at 37°C using TRIzol Max Bacterial RNA Isolation Kit (Invitrogen). RNA was further purified using the RNeasy kit (Qiagen), and quality of RNA was analyzed by Agilent 2100 BioAnalyzer (Agilent Technologies). The purified RNA was then converted to either single-stranded cDNA for use in RT-qPCR or double-stranded cDNA for use in DNA microarray analysis.

For RT-qPCR, single-stranded cDNA was synthesized from 10 µg of RNA using 100 pM random hexamer primer (Integrated DNA Technologies) and M-MuLV Reverse Transcriptase (New England Biolabs). After reverse transcription, cDNA was treated with 5 units of RNaseH (New England Biolabs) for 20 min at 37°C followed by purification with the QIAquick PCR purification kit (Qiagen). To quantify the RNA transcripts, 300 nM of primer specific to the gene of interest and 20 ng of cDNA was added to SYBR Green PCR Master Mix (Applied Biosystems) in a 20 μl reaction. Primers were designed per manufacturer’s protocol included with SYBR Green PCR Master Mix and RT-qPCR reagents. Critical threshold (Ct) values were determined using ABI Prism 7900HT Sequence Detection System (Applied Biosystems). The relative quantification of target transcripts was calculated according to the 2^-ΔΔCT^ method (58) using *ftsL* and *purC* as the endogenous control genes. Briefly, changes in Ct value (ΔCt) for the gene of interest were calculated by subtracting that gene’s average Ct from the average Ct for the endogenous control gene. The ΔCt for the mutant was then subtracted from the wild type strain’s ΔCt value to give the ΔΔCt value. Each PCR reaction was performed in triplicate and fold changes in transcript levels along with standard deviations were calculated from at least two experiments (n≥2).

For microarray analysis, Invitrogen Superscript Double-Stranded cDNA Synthesis Kit was used to generate double stranded cDNA per manufacturer’s instruction. Single-stranded cDNA was synthesized from 10 µg of RNA using 100 pM random hexamer primer (Integrated DNA Technologies) and Superscript II reverse transcriptase. Second strand synthesis was performed per manufacturer’s instructions and reaction was stopped with 0.5 M EDTA. RNA was then digested using RNaseA (25 μg/ml final concentration) followed by treatment with phenol:chloroform and precipitation with ethanol. Double-stranded cDNA was further purified with the QIAquick PCR purification kit (Qiagen) and quality tested by BioAanalyzer. Cy3 fluorescently labeled cDNA was used to probe array slides printed with 4,254 *E. coli* ORFs. Array slides contained 8 probes per gene (in duplicate) corresponding to roughly 72,000 probes per sample. Sample labeling with Cy3 fluorescent dye, hybridization to the 4-plex array (0771112 *E. coli* K-12 EXP X4, Catalog number A6697-00-01), washing, and one-color scanning was performed by Roche Nimblegen in accordance with their standard protocol. Analysis of gene expression profiles was performed using ArrayStar 2.0 software (DNASTAR) with a focus on genes with ≥2-fold change in gene expression. P-values were generated with the Student’s t-Test and false positives were minimized with FDR (false discovery rate) analysis (59).

### Enzymatic assays

The β-galactosidase assay was done following the Miller method (60). Assays were carried out with at least two independent cultures. The β-galactosidase activity was expressed as Miller Units (MU; 60). In some instances, kinetic analysis of enzyme activity was carried out using a VersaMax (Molecular Dynamics) microtiter plate reader in quadruplicate and activity was measured as the rate of ONPG cleavage divided by the cell density in each well.

### Electron paramagnetic resonance (EPR) spectroscopy

Free iron concentration in whole cells was determined by EPR spectroscopic analysis (32) with some modifications. Briefly, overnight grown bacterial cultures were diluted 1:100 in 200 ml of LB and grown shaking at 37°C until OD_600_ of 0.8. Cells were pelleted by centrifugation in a GSA rotor (Sorvall) for 10 min at 6,000 x *g*. Pellets were resuspended in 10 ml of LB containing 10 mM DTPA (to chelate extracellular iron) and 20 mM desferrioxamine (to chelate intracellular free or accessible ferric iron) and incubated shaking for 37°C for 15 min. Cells were pelleted as before and washed twice with 5 ml of ice-cold 20 mM Tris-HCl, pH 7.4. The final cell pellet was resuspended in 0.3 ml of ice cold 20 mM Tris-HCl, pH 7.4, containing 30% glycerol. A 250 μl aliquot of this cell suspension was placed in a quartz EPR tube (length: 250 mm, external diameter: 4 mm; Wilmad-Labglass). Tubes were frozen in loosely packed dry ice and then transferred to -80°C until the EPR analysis. The remaining cells were diluted 10^3^-fold to determine OD_600_. Iron standards were prepared from a freshly prepared 10 mM FeCl_3_.6H_2_O stock in a buffer containing 20 mM Tris-HCl, pH 7.4, and 1 mM desferrioxamine. Theoretical concentrations of iron standards were 100, 50, 25, 10, 5, and 0 μM. The actual iron concentrations were determined by measuring OD_420_ of each standard and using the formula: molar concentration = A_420_/ε where [Fe]ε is 2.865 mM^-1^ Cm^-1^. A 250 μl aliquot of each standard was placed in separate EPR tubes that were then frozen. The standard curve was generated by plotting EPR signals against actual iron concentrations (Fig. S6). Free iron concentration for each strain was determined from the EPR data and the standard curve. Intracellular free iron concentration was then deduced by integrating intracellular volume of the cell (1 ml of 1.0 OD_600_ cells has an intracellular volume of 0.00052 ml; Jim Imlay, personal communication) and using the formula: intracellular free iron concentration = [Fe] from standard curve/cell paste OD_600_ x 0.00052 ml.

EPR measurements were carried out at the EPR Facility at Arizona State University. Continuous wave EPR spectra were recorded using an ELEXSYS E580 CW X-band spectrometer (Bruker, Rheinstetten, Germany) equipped with a Model 900 EPL liquid helium cryostat (Oxford Instruments, Oxfordshire, UK). For all measurements, the magnetic field modulation frequency was 100 kHz, the amplitude was 1.25 mT, the microwave power was 10 mW, the microwave frequency was 9.44 GHz, the sweep time was 42 s, and the temperature was 20 K.

### OmpR purification

OmpR was purified from BL21(DE3) cultures carrying a pET24-*ompR*_*6His*_ plasmid. Overnight cultures, grown without IPTG, were diluted 1:100 in 1 L LB and grown with vigorous shaking for 90 min and then supplemented with IPTG and grown for another 2 h. Cells were pelleted, washed with 10 mM Tris-HCl pH 7.5, resuspended in lysis buffer (10 mM Tris-HCl pH 7.5, 1 mM EDTA, 100 µg/ml lysozyme) and incubated on ice for 30 min. MgCl_2_ (10 mM final), PMSF (2 mM final) and DNase I (40 µg/ml final) were then added to the cell suspension. Cells were lysed by passing through the French pressure cell three times and the lysate was centrifuged at low speed to remove unlysed cells. Envelopes were removed from the lysate by ultracentrifugation at 105,000 x *g* for an hour at 4°C. Supernatant was filtered through a 0.45 µM syringe filter and the filtrate was subjected to nickel affinity column chromatography using buffers for protein binding (20 mM sodium phosphate [pH 7.4], 20 mM imidazole and 50 mM NaCl), washing (20 mM sodium phosphate [pH 7.4], 50 mM imidazole and 300 mM NaCl) and elution (20 mM sodium phosphate [pH 7.4], 300 mM imidazole and 300 mM NaCl). Samples from eluted fractions were analyzed by SDS-PAGE and protein bands were visualized after Coomassie blue staining (Fig. S3). Fractions representing OmpR_6His_ peaks were pooled and dialyzed against a buffer containing 20 mM sodium phosphate (pH 7.4) and 300 mM NaCl. Purified proteins were stored at 4°C in the dialysis buffer supplemented with glycerol (5% final), EDTA (0.1 mM final) and DTT (0.1 mM final).

### Electrophoretic mobility gel shift assays (EMSA)

EMSA was carried out using the LightShift Chemiluminescent EMSA Kit (Thermo Scientific). *ompC* and *feoABC* promoter fragments were generated by PCR using primers specific to the region of interest, with one of the primers biotinylated. Biotin-labeled DNA probes (20 fmol), purified OmpR_6His_ (100 pmol) and other relevant reagents provided with the kit were incubated for 20 min at room temperature, and the reaction was stopped by adding 5x loading buffer. The mixture was analyzed by 5% acrylamide gel, electro-blotted onto PVDF Immobilon-P membrane (Millipore) using a Mini Trans-Blot Cell (Bio-Rad). After transfer, DNA was cross-linked to the membrane using Hoefer Ultraviolet Crosslinker and incubated with Stabilized Streptavidin-HRP Conjugate for an hour. DNA was detected by the Molecular Imager ChemiDoc XRS system (Bio-Rad) after incubating the membrane for 5 min with freshly mixed Luminol/Enhancer and Stable Peroxide solutions.

## Acknowledgments

This work was supported in part by grants from the NIH (GM048167 and AI117150, both now completed) and the School of Life Sciences, Arizona State University. We are indebted to Ananya Sen and Yidan Zhou in Jim Imlay’s lab for training RM in the EPR analysis. The initial EPR work at the University of Illinois, Urbana-Champaign facility was supported by GM49640 to JI. The Final EPR analysis was conducted at the Arizona State University EPR facility. We are grateful to Dr. Marco Flores for his assistance in the EPR analysis at ASU and constructing Figure 3.

## Supplementary Figure Legend (Gerken et al.)

**Figure S1**. Determination of *fepA, fhuA*, and *feo* expression using chromosomally integrated *lacZ* fusions. β-galactosidase activities (shown in Miller units) were measured from at three independent biological replicates in various genetic background as shown. Error bars represent standard deviation.

**Figure S2**. Effects of OmrA and OmrB deletion on *fecA* and *fepA* in wild type and EnvZ_R397L_ backgrounds. Determination of *fecA* and *fepA* expression was carried out by real-time quantitative PCR (RT-qPCR). Data were obtained from two independent biological samples and two technical replicates. Error bars represent standard deviation.

**Figure S3**. SDS-PAGE analysis of fractions containing OmpR_His_ obtained after nickel affinity chromatography. Protein samples were visualized after Coomassie Brilliant Blue staining. Fractions shown in the box were pooled and used for DNA binding studies. The position of OmpR_His_ is shown.

**Figure S4**. Determination of *ompC*::*lacZ* activities of *constructs* carrying two different lengths of the *ompC* promoter regions in front of a promoter-less *lacZ* gene. *ompC145*::*lacZ* and *ompC250*::*lacZ* contain 145 and 250 bp, respectively, of the *ompC* promoter region including ATG. The β-galactosidase activities, in *ompR*^+^ or Δ*ompR* backgrounds, were measured from two independent cultures grown to late log phase. Error bars represent standard deviation.

**Figure S5**. Effects of Δ*ompC* and Δ*ompF* gene mutations on the growth of wild type and Δ*aroB* strains. Bacterial growth was monitored on an LBA plate incubated at 37°C for 24 h. The relevant genotypes are shown.

**Figure S6**. Standard curve of ferric chloride solutions of known concentrations. The actual concentrations of ferric chloride solutions were determined by measuring absorbance at 420 nm and then using the formula: molar concentration = A_420_/ε where [Fe]ε is 2.865 mM^-1^ Cm^-1^. These values were plotted against the values obtained for the same solutions by electron paramagnetic resonance (EPR) spectroscopy. We used peak-to-peak EPR measurements instead of the double integration values due to a greater confidence of the former at lower ferric chloride concentrations. The standard curve was used to measure iron concentrations in the whole cell.

